# DyME: An MD-based engine exploiting HTP mutagenesis for protein engineering and recognition mimicry

**DOI:** 10.64898/2026.04.10.717642

**Authors:** Pedro M. Guillem-Gloria, Gloria Ruiz-Gómez, M. Teresa Pisabarro

## Abstract

Protein recognition mimicry is of great interest in the field of molecular bioengineering and rational design, with mutagenesis frequently employed to analyze the effects of altering amino acids involved in molecular recognition. The conformational and energetic effects of such alterations can be investigated in detail with the help of molecular dynamics (MD) methodologies. While existing MD-based computational tools can be used to explore a particular set of mutations at a time, suitable for small-scale studies, high-throughput (HTP) exploration of protein recognition for engineering purposes would greatly benefit from an integrative platform that streamlines preparation, mutagenesis, simulation and post-processing of up to several thousand molecular systems, along with robust tools for comprehensive and straightforward comparative analysis.

DyME (Dynamic Mutagenesis Engine) is a distributed platform that enables systematic investigations of protein recognition mimicry by combining HTP mutagenesis, solvated MD simulations and a Toolbox for comparative analysis (TCA), including interfacial water-site mapping. DyME uses 3D structural information of any protein-protein or protein-DNA complex as input. Its automated MD-based mutagenesis engine facilitates systematic investigation of how site-specific alterations affect recognition, enabling the organization of single, double and triple modifications into combinatorial libraries for comprehensive comparative analysis. In DyME, relevant MD trajectory-derived data is scavenged and stored into a central database, providing aggregation capabilities that ease multi-feature analysis across an extensive collection of simulations. An interactive web-GUI and specialized widgets simplify preparation and efficient molecular and numerical comparative exploration. DyME’s capabilities are evaluated using available experimental data. Its source code is available at https://github.com/pisabarro-group/DYME

## Introduction

In protein recognition, interfacial amino acids play deterministic roles [1]. Their conformational and functional characteristics can be altered to gain binding stability and selectivity, engineer new variants of the interacting proteins with customized functionalities, or designing mimetics [2]. Although systematic interfacial amino acids alteration is a powerful strategy to adjust key properties of proteins [3] [4], static structure-based methods (SSBM) are limited in their ability to account for the complexity of the dynamic mechanisms governing biological systems. In such scenario, molecular dynamics (MD) is considered the golden standard [5] [6], as the emergence of graphical processing units (GPUs) as efficient co-processors has enabled simulating large macromolecular systems at significantly reduced execution times, thus making large-scale studies feasible [7].

The combined use of mutagenesis and MD has become a standard approach in small-scale studies, serving various applications of rational design such as engineering of antibodies [8], affinity [9], stability [10] and lead optimization [6] [11]. In such studies, each MD simulation typically corresponds to a single set of mutations. However, despite recent advances in computational resources, exploring protein mimicry through large-scale interfacial mutagenesis remains challenging due to the complexity of conformational dynamics in protein interactions.

Combining and interpreting MD trajectory-data from large mutational sets is nearly infeasible without advanced multi-feature analysis tools. However, most existing computational tools are designed to handle simulations individually. To address tackle this limitation, efficient automated software is needed to extract, compare and summarize design-relevant insights from large datasets, enabling high-content comprehensive exploration of protein recognition. In addition, tracking of water molecules represents a crucial aspect to take into account in PRM studies, as interfacial water-mediated interactions are essential participants of molecular recognition and function [12] [13].

This scenario presents an opportunity for innovation in terms of handling, production and processing of large collections of MD simulation data. At such scales, the need for modern data management paradigms becomes evident. Document-based database engines can accommodate these needs [14]. Recent initiatives like the MDDB project have identified and implemented notable improvements based on database-driven storage of MD simulations and MD feature data, consolidating a collaborative repository for collective meta-analysis [15] [16]. A data paradigm based on similar principles could be exploited to address unmet requirements for MD-based HTP mutagenesis in terms of data homogenization, normalization, federation and complex query capabilities, thus accelerating efficient comparative analysis of multidimensional data across many simulations.

Up to date, a small number of workflows that automate HTP mutagenesis have been reported. For instance, a semi-automatic pipeline that explores saturation mutagenesis built by Chiappori et. al. in 2009 [17] using Perl [18], Modeller [19], Autodock [20] and FoldX [21]. FoldX probes singlet and doublet permutations at a receptor in complex with its ligand. It runs site-specific ligand docking for each receptor permutation and uses the FoldX stability function to identify favorable energy gains. A similar workflow called MutateX [22] provides an automation layer to FoldX for HTP mutagenesis exploration. Both tools use energy-minimized structures instead of MD simulations, therefore remaining constrained to the above-mentioned limitations of SSBM. A more recent tool, MDFit [23], can be provided a protein structure and a library of pre-docked mutated ligands as input, followed by automated MD simulations using Desmond [24]. However, this tool doesn’t perform automated mutagenesis, and its ligand library must be built manually. Other tools like StreaMD [25] can automate MD simulations when provided a library of existing input structures and assist post-processing using GROMACS [33, 34]. While these tools cover some of the steps to make large-scale MD-based studies feasible, an expert tool covering a broader scope and all steps in a single workflow has not been reported.

In response to these needs, we have created DyME; a specialized platform that bundles HTP mutagenesis, MD simulations, feature scavenging and a GUI toolbox for extensive comparative analysis. DyME’s workflow covers all steps to seamlessly generate, simulate and explore up to thousands of mutant permutations. It uses Amber [28] file formats for standardization, Modeller [19] for structural mutagenesis, OpenMM [29] for automating MD simulations, MongoDB for efficient data storage, distributed agent nodes for task parallelization, and orchestrates several existing tools and libraries like ParmED [30], MDTraj [31], MDAnalysis [32], CPPTRAJ/PYTRAJ [33] [34], MMPBSA.py [35] and AmberTools [28] for post-processing. Furthermore, DyME implements a powerful new water-site mapping analysis module (to be published) and conveniently supports mutable molecular objects with non-standard constituents, like synthetic amino acids, thereby advancing applications in mimicry design.

DyME is built using high-level programming languages (Python, Shellscript, Javascript, PHP) and other state-of-the-art server-side technologies. It incorporates a web-based frontend and a distributed backend, which communicate through an API. Its workflow execution uses the producer-consumer strategy implemented asynchronously through processing nodes. Events are propagated with status codes using periodic database polling. The DyME system is based on Linux and bundled on Docker/Apptainer containers for ease of distribution.

We provide a case study showcasing the utility and validation of DyME in preparation and execution of a MD-based mutagenesis study for affinity and specificity design in protein mimicry using a experimentally validated data-set [36]. Numerical comparative analysis revealed correlation between the binding free energies (ΔG) obtained within DyME and the reported experimental *K*_d_ values. Furthermore, DyME’s water-site mapping was able to elucidate water-mediated interactions identified experimentally [37–39], demonstrating DyME’s capabilities and highlighting its potential for assisting rational engineering.

## Design and Implementation

### Workflow and Concept Development

In the DyME workflow (Fig 1), interfacial residues (or “anchor points”) within a protein complex structure are specified during an input data preparation stage (Fig 1A), subsequently undergoing adjustable iterative mutagenesis, followed by automated MD simulations (Fig 1B) and feature scavenging routines (Fig 1C). Output data is collected and stored in a central database (Fig 1D) and efficiently aggregated (Fig 1E) to serve responsive UI widgets, providing visual representations and numerical information for comparative analysis on-demand (Fig 1F).

**Fig 1.**
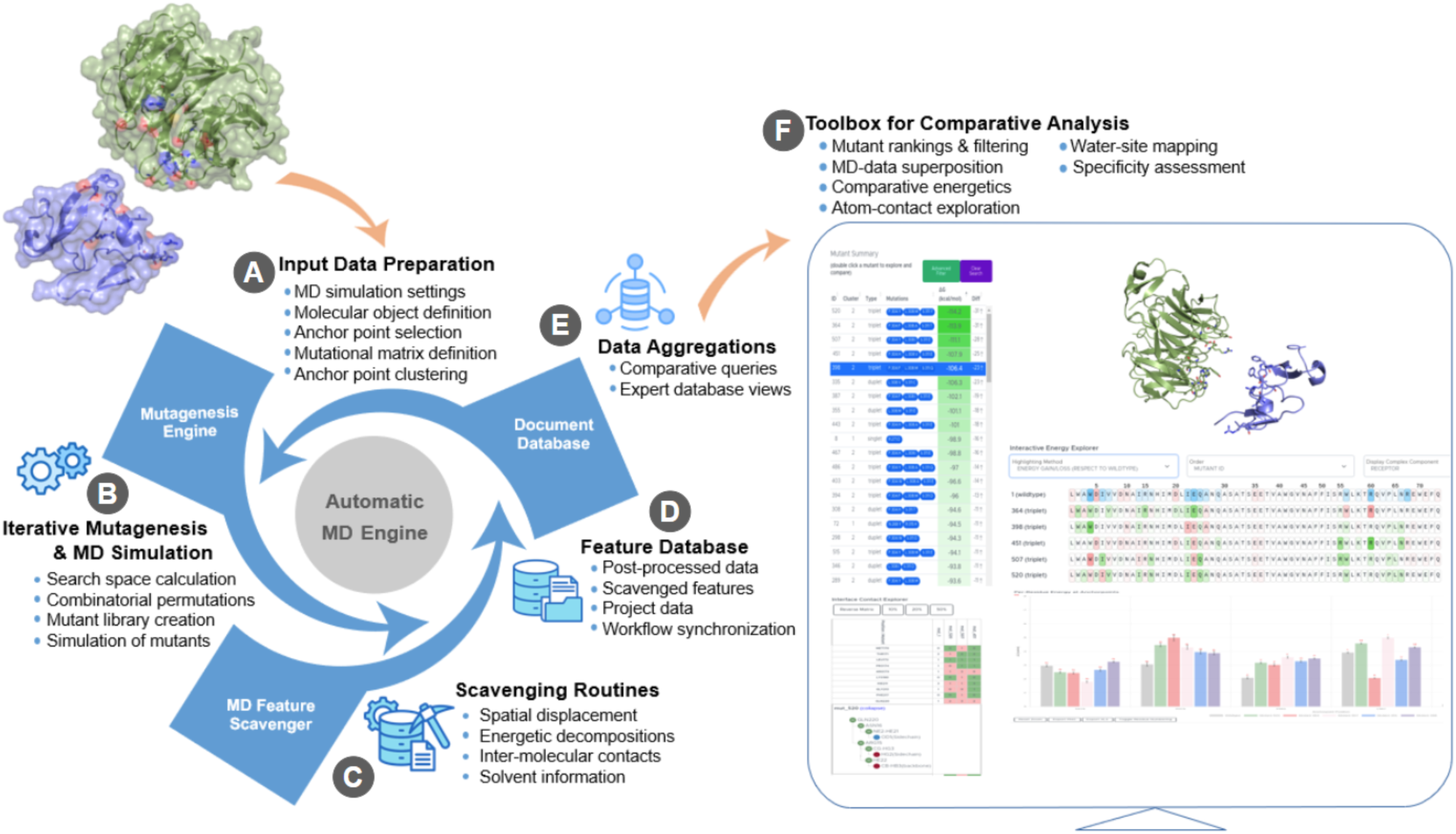
DyME Workflow. **(A) Input data preparation.** The 3D coordinates of a protein in complex with another protein, a peptide or DNA molecule are provided as the *wild-type* (WT) conformation. Each molecule in the WT complex is defined as a *“molecular object”*, and one such object is designated by the user as “*mutable*”. DyME analyzes inter-molecular contacts and presents a tentative list of *anchor points*. These are interfacial residue positions at the *“mutable object”* that qualify for mutagenesis. The user is given the option to modify DyME’s selection and configure any other position of interest. This is especially helpful for investigating positions considered relevant for analysis, even if they aren’t ultimately selected for mutation. At this stage, the user defines the settings relevant for the workflow (*i.e.* MD settings, scavenging parameters, analysis criteria). (**B**) **Iterative Mutagenesis and MD simulation engine**. The *“MD engine”* of DyME takes care of the preparation and subsequent simulation of each mutant in the library. DyME automates the creation of MD input files, including 3D structure generation of each mutant in a distributed on-demand fashion, with tasks allocated across (one or multiple) MD simulation nodes. The *MD engine* is designed to run simulations asynchronously on idle GPU hardware. Each node produces the MD inputs and the corresponding MD trajectory output for every mutant assigned. To process mutants, MD nodes periodically poll for pending entries in the library awaiting generation and simulation. (**C**) **Scavenging routines.** Upon completion of each simulation, *“scavenging routines”* in DyME automatically extract MD trajectory information (such as RMSDs, binding free energies, pairwise/per-residue energy decompositions, inter-molecular contact frequencies and solvent information) to be stored in a central *“Feature Database”* (Fig 1D), indexed for efficient aggregation and subsequent analysis (Fig 1E-F). All assets necessary for trajectory post-processing are created and managed internally. A Scavenging node designed for this purpose produces data asynchronously on a first-come-first-served basis. A major advantage of this approach is that exploration of partial results is possible as soon as data becomes available. (**D**) **Feature Database.** DyME uses database collections to store project-specific data and curated MD-data. This database also serves as a central hub for synchronization among different units of the workflow. To facilitate synchronization, mutant entries at the database contain status code that determines the processing task being performed and which should be performed next (*i.e.* simulating or scavenging). **(E) Data Aggregations.** DyME exploits native query aggregation capabilities at the database to enable high-speed correlational queries among millions of records containing features of several MD simulations. This is useful for summarizing, grouping or transforming data at unprecedented speed and simplicity. (**F**) **Toolbox for Comparative Analysis (TCA).** An interactive web-based GUI provides DyME with a set of comprehensive interfaces for investigation of MD-derived data, generated from query aggregations. This facilitates instantaneous overlapping of features from several MD simulations of interest, enabling straightforward comparative analysis and expedited identification of relevant features for engineering purposes (*e.g.* identification of best mutations for affinity or selectivity optimization).

A critical requirement for HTP mutagenesis is generating a collection of mutated 3D structures (comprising one or several point mutations) constituting a “*mutant library”*. DyME uses the provided WT structure as template and automates mutagenesis by collecting a matrix of selected residue substitutions at each *anchor point*. This mutants-matrix is used at a later stage to calculate all possible mutant permutations, and a database collection stores the “*mutant library”*. DyME incorporates “*anchor point clustering”* capabilities to deal with: *i)* groupings of mutable anchor points of interest distant in 3D space, and *ii)* different groupings of adjacent mutable positions. The mutagenesis of *anchor points* can be assessed individually, or collectively by grouping positions in a cluster. DyME will calculate all mutant permutations at each cluster in a singlet, doublet and triplet fashion. Each entry in the *“mutant library”* will represent a unique mutant that must undergo structural generation, preparation for MD simulation, feature scavenging and post-processing.

At a mouse-click, DyME allows verifications (*i.e.* running additional MD replicas), arbitrary mutant permutations, and generation of VMD [40] scripts with automated and pre-defined molecular representations to expedite visual trajectory analysis.

Two particularly helpful exploration tools incorporated in the DyME platform as powerful assets for investigating protein interactions for PRM are “the *“Water-site Explorer”* and the “*Specificity Finder”*. The *“Water-site Explorer”* facilitates in-depth characterization of solvent involved in recognition. At just one mouse-click, the “*Specificity Finder*” enables straight forward: *i)* comparison of binding free energies obtained from two DyME projects sharing the same *“mutable object”* (or ligand) in complex with a different *“non-mutable object”* (or receptor) in each case, and *ii)* identification of mutations (single or combinations) simultaneously resulting in increased versus decreased affinity for either protein receptor.

A DyME cycle begins when an “*exploration project*” is initialized. Each project is a dedicated namespace containing all relevant assets, including the initial WT complex, a collection of input files, simulation settings, MD outputs and their corresponding post-processed data. The generation and management of these assets at each stage is handled automatically by the workflow, requiring minimal user intervention. After finalizing all processing stages (Fig 1 A-E), users can perform comparative exploration with the TCA (Fig 1F).

### System Architecture

DyME was designed as a distributed computing platform made of several components, including a GUI, a main node, worker nodes and two storage facilities (raw disk and a central database), as outlined in Fig 2.

**Fig 2.**
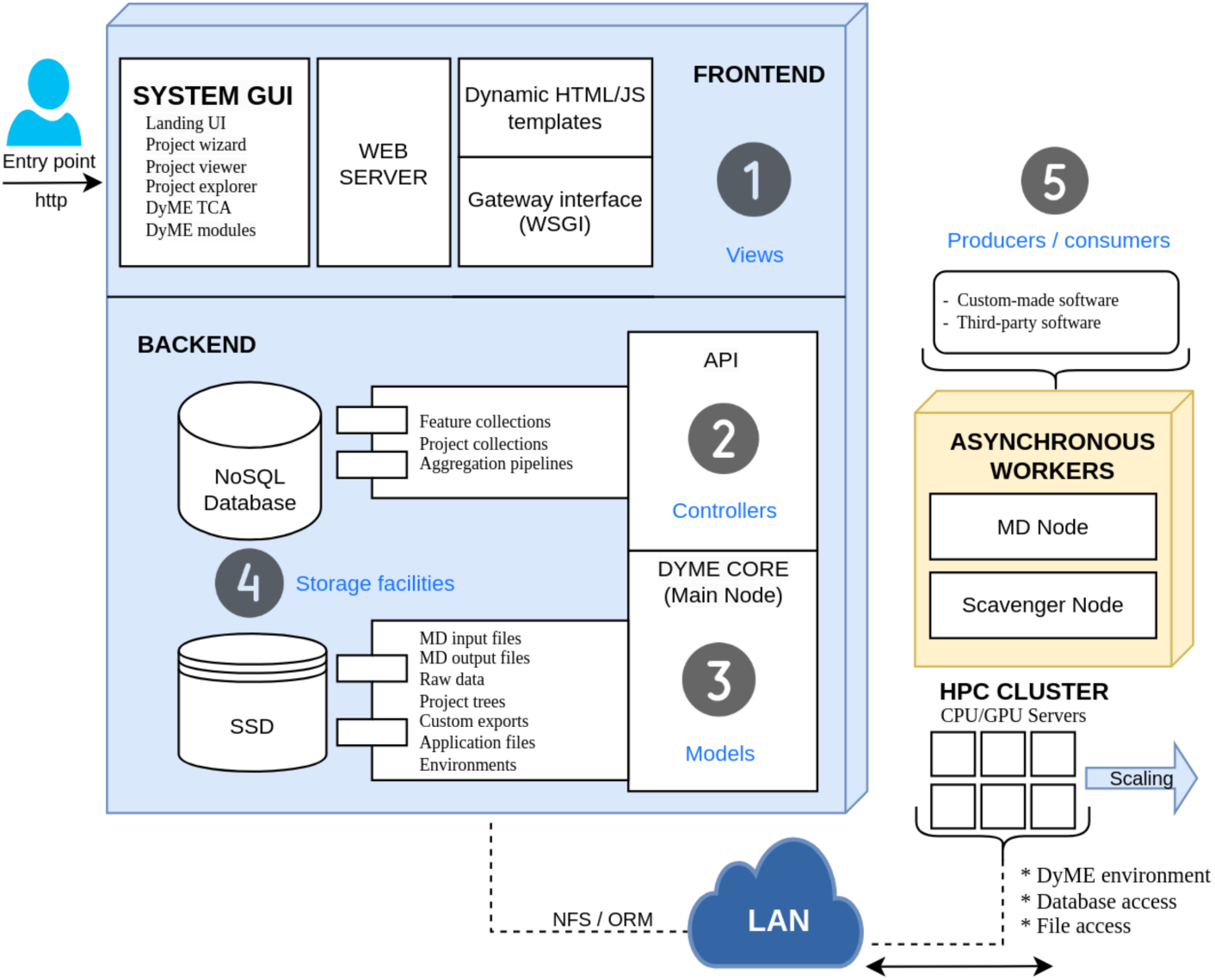
DyME system architecture. The system components (1-5) are divided into two logical groups: Frontend (1) and Backend (2-5), which communicate through a Restful API. At the backend, the main node acts as orchestrator, and worker nodes perform workflow tasks asynchronously. The software architecture resembles the Model View Controller (MVC) design paradigm. The *Controller* (business logic) layer is implemented within the backend. *Views* are served as web-based content at the front end, while *Models* and other data abstractions are handled through a database access library. Workers are individual units that scale horizontally and perform tasks for different stages of the workflow.

### Frontend Layer

The Frontend of DyME (Fig 2.1) is a web application built with HTML, PHP, JS, Bootstrap, JQuery and Ajax. The entry point is a single PHP index file. The GUI sections are divided into smaller modules, each designed as a pair of PHP/JS files. Widgets and components of the GUI consume the DyME API via asynchronous Ajax calls. Responses are received in JSON format and handled by the component that originated the request.

### Backend Layer

Backend components (API, asynchronous workers and main node) are built in Python 3.11. The API (Fig 2.2) is a Flask application exposed through a WSGI gateway using a legacy web server (Apache). The gateway uses a single Anaconda container where libraries, dependencies and third-party tools reside (see

Table A on S1 Appendix). Methods that communicate with the Frontend reside in the API. Base instances (*i.e.* main node) and other components (*i.e.* asynchronous workers) are part of the DyME core and communicate through active database polling (Fig 2.3).

**Fig 3.**
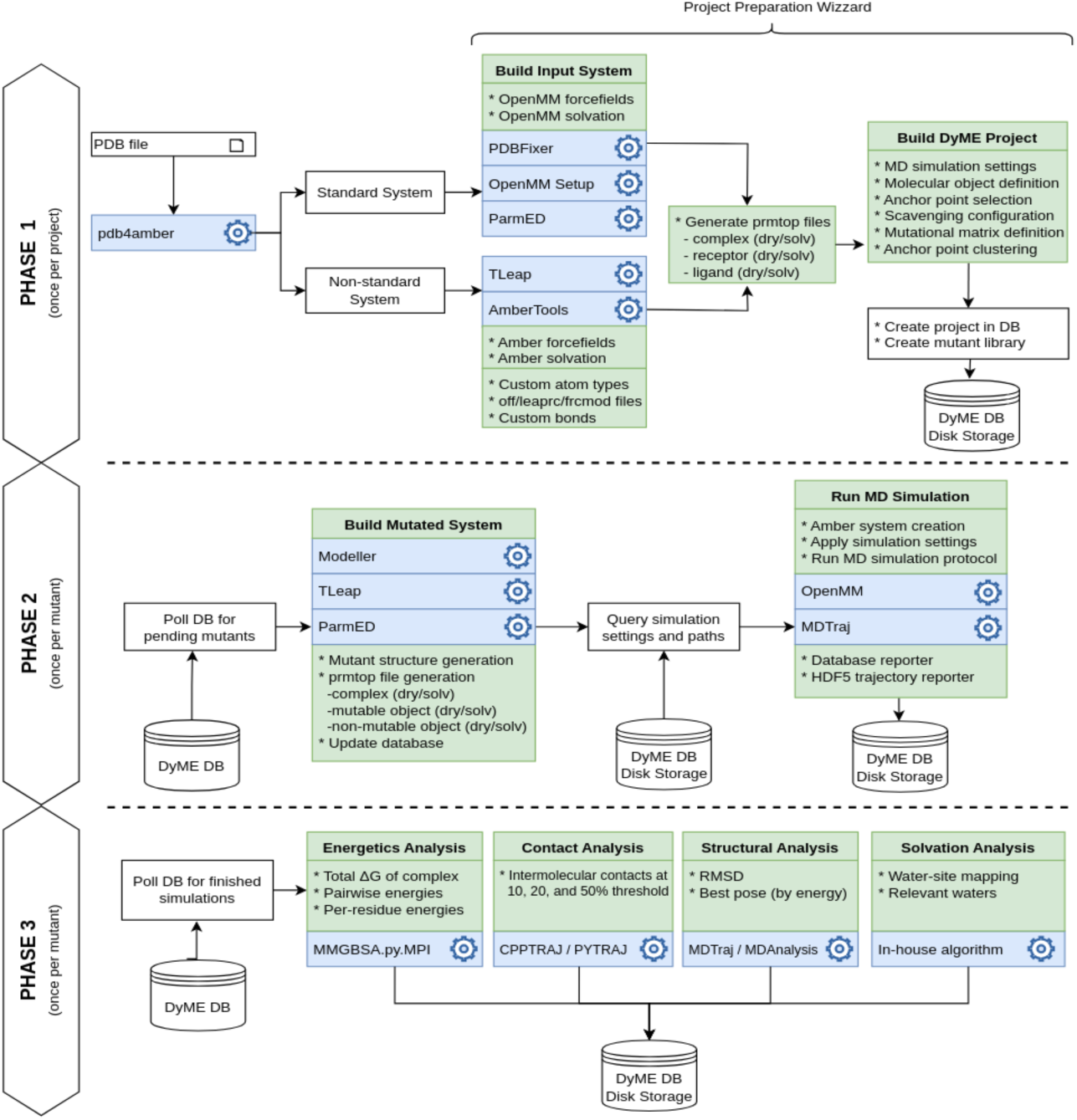
Implementation overview of the DyME workflow. The corresponding phase is indicated at the left. Elements in blue denote the name of the tool used to perform tasks. Elements in green denote actions, tasks, output products or resources.

### Main Node

This entity contains the database engine, the web server, the core components of DyME, and the necessary abstractions for business logic execution at the backend. This node runs as an individual container.

### Worker Nodes (Asynchronous Workers)

The workflow processing subsystem is built into two “*worker nodes”* (Fig 2.5). Each worker node embeds routines specific to a set of tasks, such as running MD simulations, scavenging data, or generating inputs for other tasks. Worker node instances can run on one or many distributed servers.

### Storage

DyME uses two storage facilities (Fig 2.4): a document-based database engine (*MongoDB*) and a dedicated disk directory (ideally a shared NSF partition). Each project in DyME is given its own directory tree. The database consists of 4 collections (“default_settings”, “projects”, “mutants”, and “processed_data”), and 3 logical views containing aggregation pipelines tailored for TCA integration (see Fig A on S1 Appendix). The database access module is implemented using a native driver (*pymongo*) and used by all components for persistent connectivity.

### Processing Paradigm

The DyME workflow follows the producer-consumer paradigm. Worker nodes initiate tasks by polling the database for pending jobs. Upon initiating or completing a task, or a sequence of operations for a given task, the “status” of the task is updated accordingly so that other nodes perform successive steps in the workflow. Nodes reside within isolated containers and execute distinct responsibilities within the distributed system. A typical DyME deployment is made of one individual main node and one -or several-instances of the MD or Scavenger nodes across servers. The platform scales horizontally by deploying worker nodes in as many distributed servers as desired.

### Deployment and Distribution concept

DyME uses containers running Ubuntu Linux. The main node container exposes ports 8080 for web GUI and API access, and 27017 for database access. The GUI can be accessed from any web browser in the local network using the hostname or IP address of the main node container as URL. Worker nodes are deployed as apptainer/singularity container instances.

## Results

### Components and Method Development

The DyME subsystem was split into three computational phases, each representing an independent stage of the workflow (Fig 3). Phase 1 handles the upload of the initial 3D structure of the molecular complex, the definition of MD settings, anchor point selection, scavenging parameters, mutant library generation and project namespace creation. Phase 2 implements mutant generation and automation of MD simulations. Phase 3 handles scavenging, feature extraction and storage of MD-derived data.

### Phase 1

Molecular systems without the need for custom bonds, custom force fields or non-standard parameters undergo the “*Standard System*” preparation path in the backend. Any other non-standard system undergoes the “*Non-standard System*” preparation path, which accepts custom Amber libraries (.lib, .frcmod and .off files). DyME uses Amber file-format conventions for standardization. Prior to uploading the initial WT coordinates, the input PDB file must be prepared with *pdb4amber* [28]. This tool removes existing water molecules and hydrogens,adds missing heavy atoms, re-numbers amino acid positions, verifies chain identifiers and performs corrective operations that enforce Amber compatibility. A key benefit of using Amber conventions is that other existing tools are compatible with its topology, coordinates and parameters (*dry* and solvated -*solv*-) formats. This offers great versatility for handling custom atom types, coordination states, force field modifications and molecular systems with non-standard constituents, like synthetic amino acids. Likewise, OpenMM [29], used for the MD simulations, conveniently supports the creation of its “*System objects*” from Amber files.

Project preparation wizard: Phase 1 implements an interactive GUI that guides the user into project creation and enforces input validation. This interface is delivered as a “*project preparation wizard*” (Fig 4 and 5), which consists of 3 steps to define project settings, MD simulation settings, molecular object definitions, scavenging parameters, mutagenesis settings and anchor point clustering.

**Fig 4.**
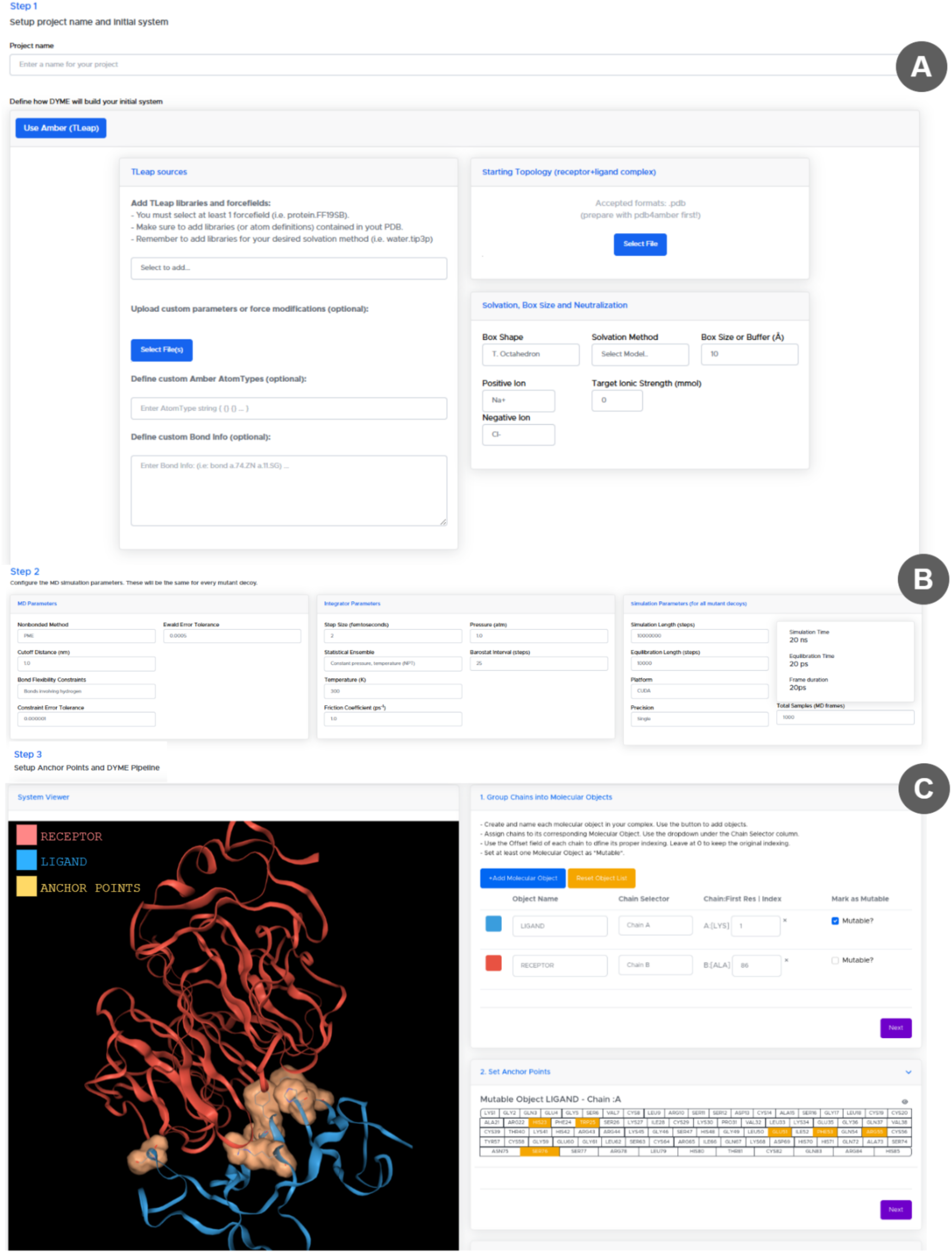
Project preparation wizard GUI. **(A) Step 1 – <u>Input PDB preparation</u>**. The 3D structure of the initial WT complex is uploaded into the backend and iterated through a PDBFixer object [29], which extracts PDB data (*i.e.* chain identifiers, residue numbers, type of atoms, etc.) and determines which options should be presented to the user during the wizard. At this stage, the system’s parameters are collected (*i.e.* solvation model, ionic neutralization strategy, box size, etc). **(B) Step 2 - <u>MD simulation parameters setup.</u>** The user configures several MD simulation settings such as periodic boundary conditions, restraints, temperature, statistical ensembles, step size, equilibration steps and total production steps. The wizard provides dynamic form elements to collect these settings appropriately, in a similar fashion than the tool OpenMM-Setup by Eastman, *et al* [29]. **(C) Step 3 - <u>Mutagenesis pipeline parameters setup</u>**. The user must define abstractions for each molecular object (*i.e.* mutable object and positions). The wizard provides tools to group different chains as a single molecular object. The *“mutable object”* is defined using a checkbox. Each object is assigned a color to ease visual inspection. NGLViewer [41,42] is used to provide a 3D interactive representation of the complex structure. This molecular visualization window is responsive to user inputs. Anchor point definition is carried out using an interactive grid of buttons containing each position in the *“mutable object”*. Hovering the mouse over a button in the grid highlights interactively the van der Waals surface of the corresponding amino acid side chain in the 3D viewer.

**Fig 5.**
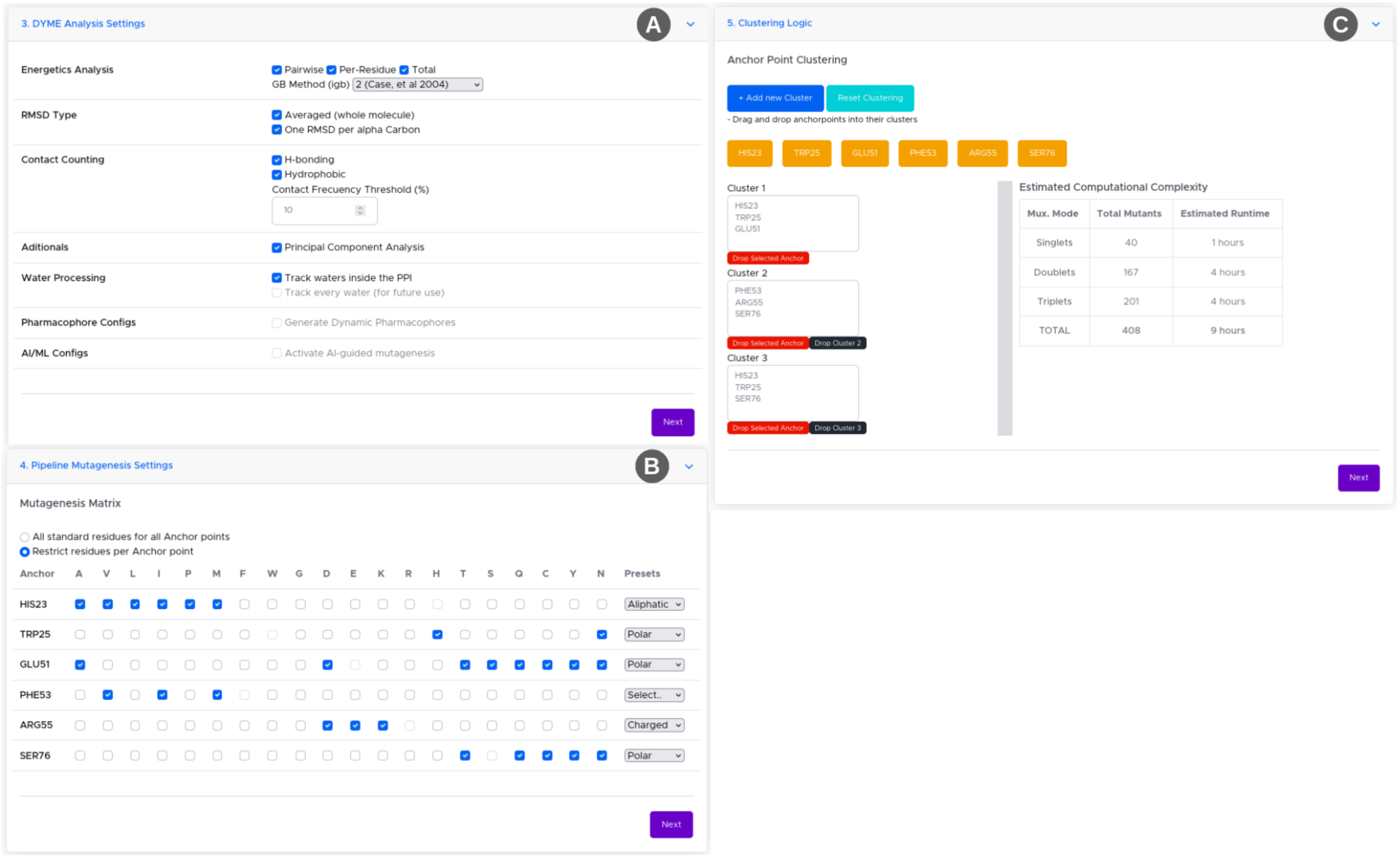
Scavenger settings, mutagenesis and clustering during project preparation wizard. **(A) Scavenger configuration.** It configures several parameters for scavenging routines (see details in Phase3). **(B) Mutagenesis settings.** It provides a grid for selecting the residue substitutions to be introduced at each anchor point. The grid is generated dynamically from the list of anchor points defined by the user. Residues can be selected one at a time or using the “Preset” combo boxes at the end of each row. **(C) Anchor point Clustering**. It provides the means for anchor point grouping. Top: the selected anchor points appear as residue-labeled orange buttons. Bottom left: selected anchor points can be manually dragged into one, or as many cluster boxes as desired. Users can define as many clusters as needed using the “+Add new cluster” button (in blue). Bottom right: Upon adding an anchor point to a cluster, the system re-calculates the number of estimated total mutant permutations (including all possible singlets, doublets or triplets) and displays the estimated computing time in GPU-hours.

Following the definition of *molecular objects* and *anchor points*, the wizard provides additional forms to configure scavenging parameters, a grid to select the desired substitutions at each anchor point, and anchor point clustering (Fig 5).

To complete the setup of a DyME project, a JSON object containing input settings, molecular objects, cluster definitions and MD simulation settings is sent to the backend, and a new project is created and assigned a unique ID into DyME. Lastly, the finalized *“mutant library”* for the project is created at the backend by computing non-redundant mutational combinations for all clusters (see S2 Appendix for details on Methods and Implementation). Mutant records are made unique by assigning an incremental ID and parent project ID. Each combination is stored as an individual database record and assigned the “pending” status, so that worker nodes can initiate the corresponding processing tasks.

### Phase 2

The logic of Phase 2 resides in the “MD node”. Instances of this node automatically generate MD-input files for any mutant assigned, run MD simulations on idle GPU cards, and periodically write simulation status updates to the DB. Fig 6 outlines the flow of this process.

**Fig 6.**
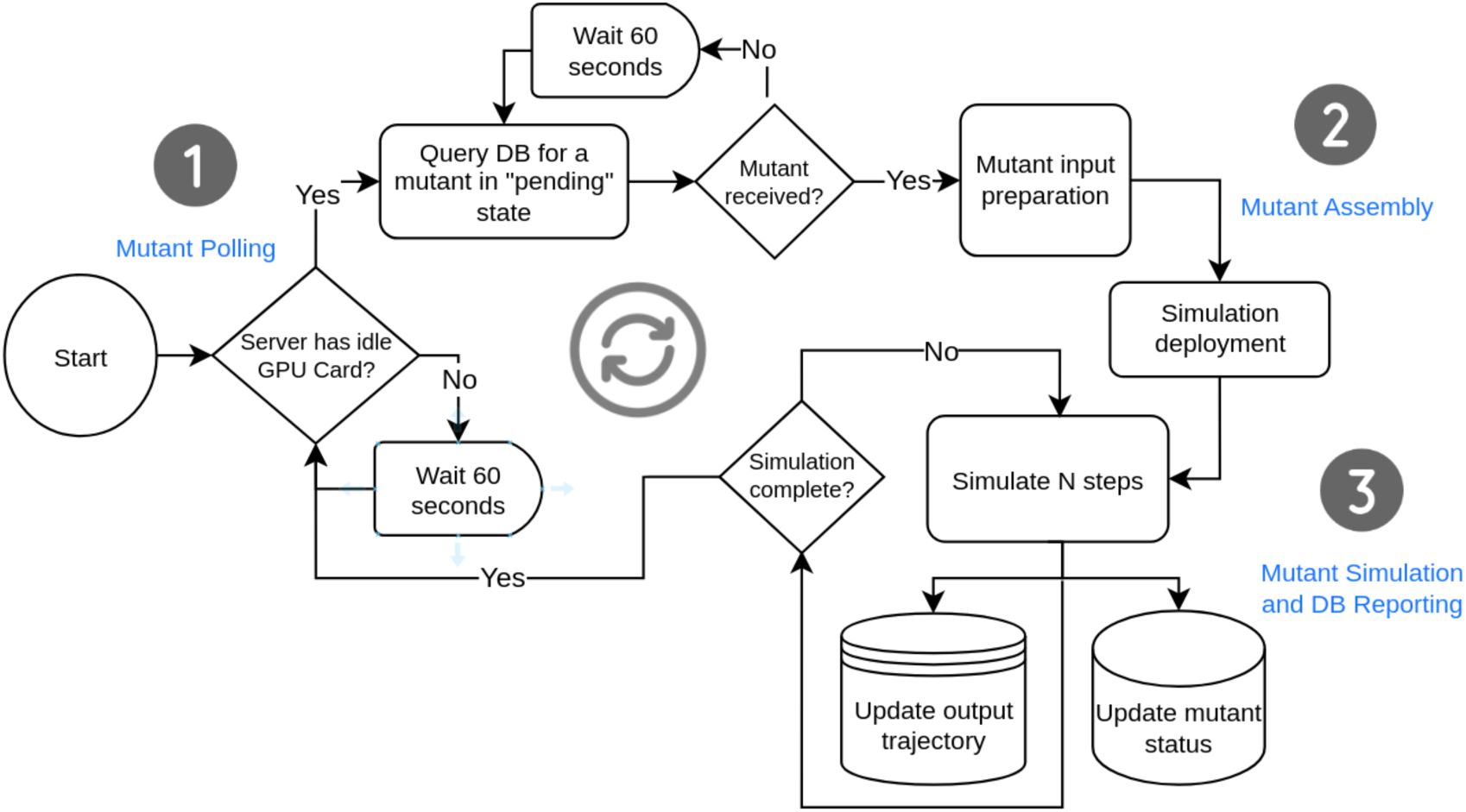
Processes and methods of the MD node, implementing Phase 2 of the DyME workflow. When initiated in a GPU server, MD nodes periodically scan the local hardware for idle GPU resources and poll the central database for mutant records on “pending” state (Fig 6.1). Both conditions are required to initiate a simulation. If a mutant is pending to be simulated, the database response delivers the project simulation settings and the necessary data to build its MD inputs. Idle GPU cards are assigned one single MD simulation at a time. The main process of the instance spawns each MD simulation as a Python subprocess (using the *multiprocessing* library).

Upon receiving the database response, the 3D structure of the pending mutant is built with the *mutate_model()* function of Modeller [19] using the initial WT structure as template (Fig 6.2). The newly created mutant structure is energy minimized within Modeller, which rotamer refinement capabilities prevent steric clashes, saved in PDB format and then loaded into Amber’s tleap tool [28] to produce additional files necessary for MD simulations. DyME embeds an automatic LEaP template generator built for this purpose, which crafts the necessary instructions to perform solvation, charge neutralization and building the corresponding Amber topologies, parameters, and coordinate files (*i.e. prmtop* and *inpcrd* files). Dry and solvated representations are created and kept for later use. The *strip()* function of MDTraj [31] is used with appropriate selector masks for separating Amber topologies into individual files (*i.e. mutable and non-mutable object*). This step guarantees that the number of atoms across file fragments remains consistent on every mutant complex, regardless of its mutations, which is required for successful energy decomposition at a later stage.

To run an MD simulation, OpenMM [29] is instantiated (Fig 6.3). The OpenMM “*System object*” is created from Amber files using the *AmberPrmtopFile()*, *AmberInpcrdFile()* and *createSystem()* functions. The OpenMM “*Context*”, “*State*” and “*Simulation*” objects are created using the pre-defined MD settings of the project. DyME uses the LangevinMiddleIntegrator [43] by default. Energy minimization is done using the OpenMM’s *minimizeEnergy()* function until convergence is achieved. The equilibration and production stages take place for the number of steps specified during project setup. At this stage, the “status” field of the mutant record in the database is updated to “processing” in order to prevent racing conditions between MD nodes.

DyME uses two OpenMM reporters: The “HDF5 Reporter” by MDTraj [31] to write trajectory frames as output, and the “DyME Database Reporter” (see S2 Appendix), which periodically updates simulation status into the database. Upon completion of an MD simulation without errors, the mutant record status field is updated to “ready_to_scavenge”. Should errors occur, the mutant record is marked as “failed”. Errors can be conveniently traced in the execution log file of each mutant process, which is stored within a mutant’s unique storage directory.

### Phase 3

The logic of Phase 3 resides in the “Scavenger node”, which implements trajectory post-processing and feature scavenging from MD simulations. The tools involved in this process include ParmED [30], MDTraj [31], MDAnalysis [32], CPPTRAJ/PYTRAJ [33,34], MMPBSA.py [35] and AmberTools [28]. Fig 7 outlines the flow chart of this process.

**Fig 7.**
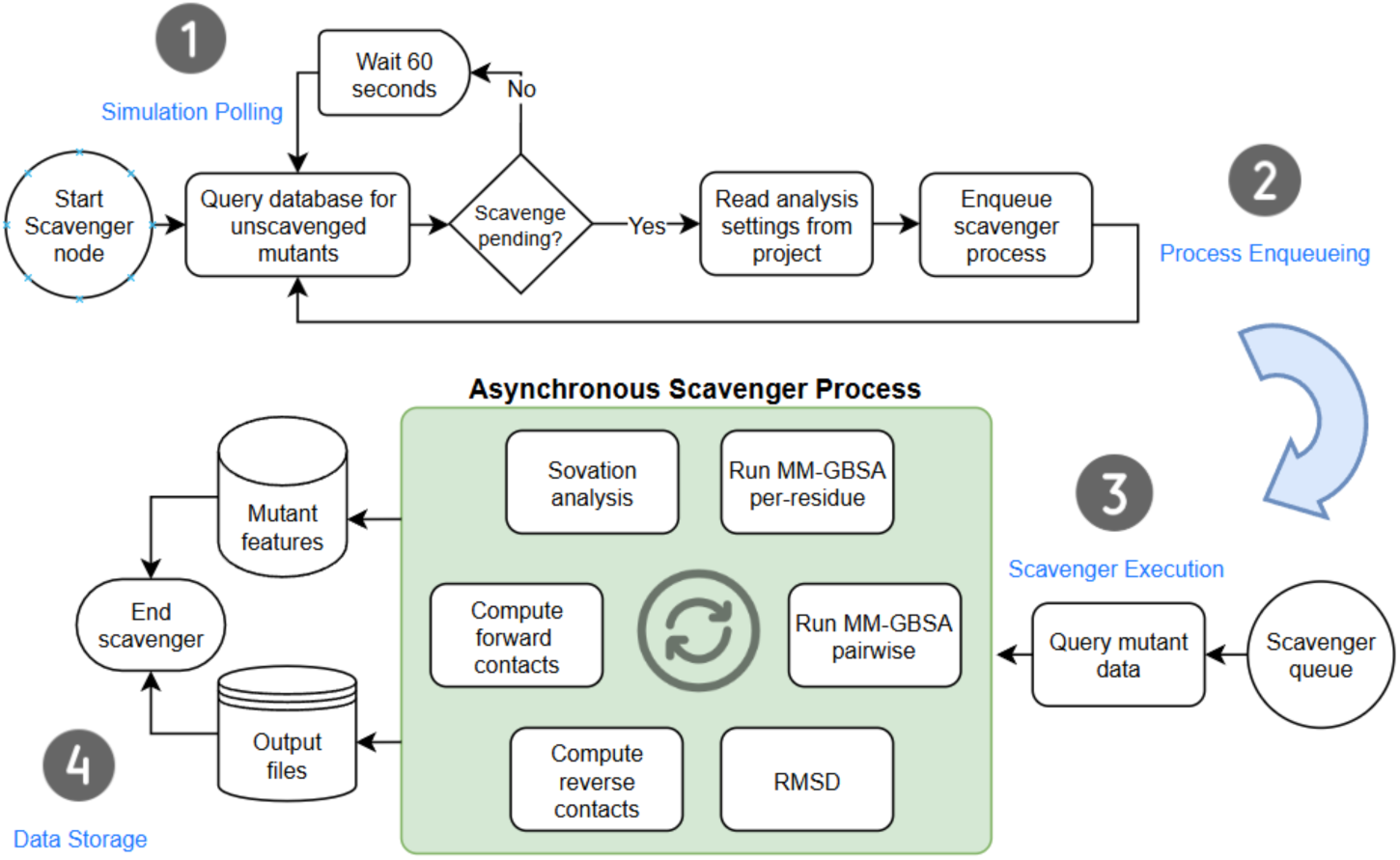
Processes and methods of the Scavenger node, implementing Phase 3 of the DyME workflow. The six tasks of the scavenging process applied after successfully completing MD simulations are highlighted in the green box.

The Scavenger node polls the database for mutant records in “*ready_to_scavenge*” status (Fig 7.1). This status code is only possible if the simulation has taken place without errors and its H5 trajectory file exists. For each mutant record returned, a *“Scavenger process”* is started. Scavenging nodes can be deployed using Slurm [44] (Fig 7.2), or manually by running the node on a server. Several mutants can undergo scavenging on a single server depending on the number of available CPU cores. The Scavenger execution (Fig 7.3) performs all tasks sequentially on every mutant, while managing MPI parallelization automatically. Upon completion of all tasks, data collections built by each task are labeled with their parent project and mutant ID and stored at the database collection “*processed_data*”, while raw output files remain available on the corresponding project directory (Fig 7.4). Computational details on collecting binding free energy calculations, atomic contact extraction, RMSD data, best frame of the trajectory and water site mapping can be found in S2 Appendix.

### Toolbox for Comparative Analysis (TCA)

DyME contains a Toolbox for Comparative Analysis (TCA) that assists users with graphical and numerical analysis of those MD simulations selected for exploration in a fully interactive fashion. A key advantage of this tool is that it allows comparisons of many simulations in parallel, which expedite visual feature recognition and rationalization. The TCA is built into two main GUI modules: the *‘Mutant Explorer’* and the *‘Specificity Finder’*.

### Mutant Explorer

This web application serves as the primary exploration resource of the TCA (Fig 8). It is composed of multiple widgets organized into the “Mutant Simulation Results” panel (Fig 8, left) and the “Data Viewer” panels (Fig 8, right).

**Fig 8.**
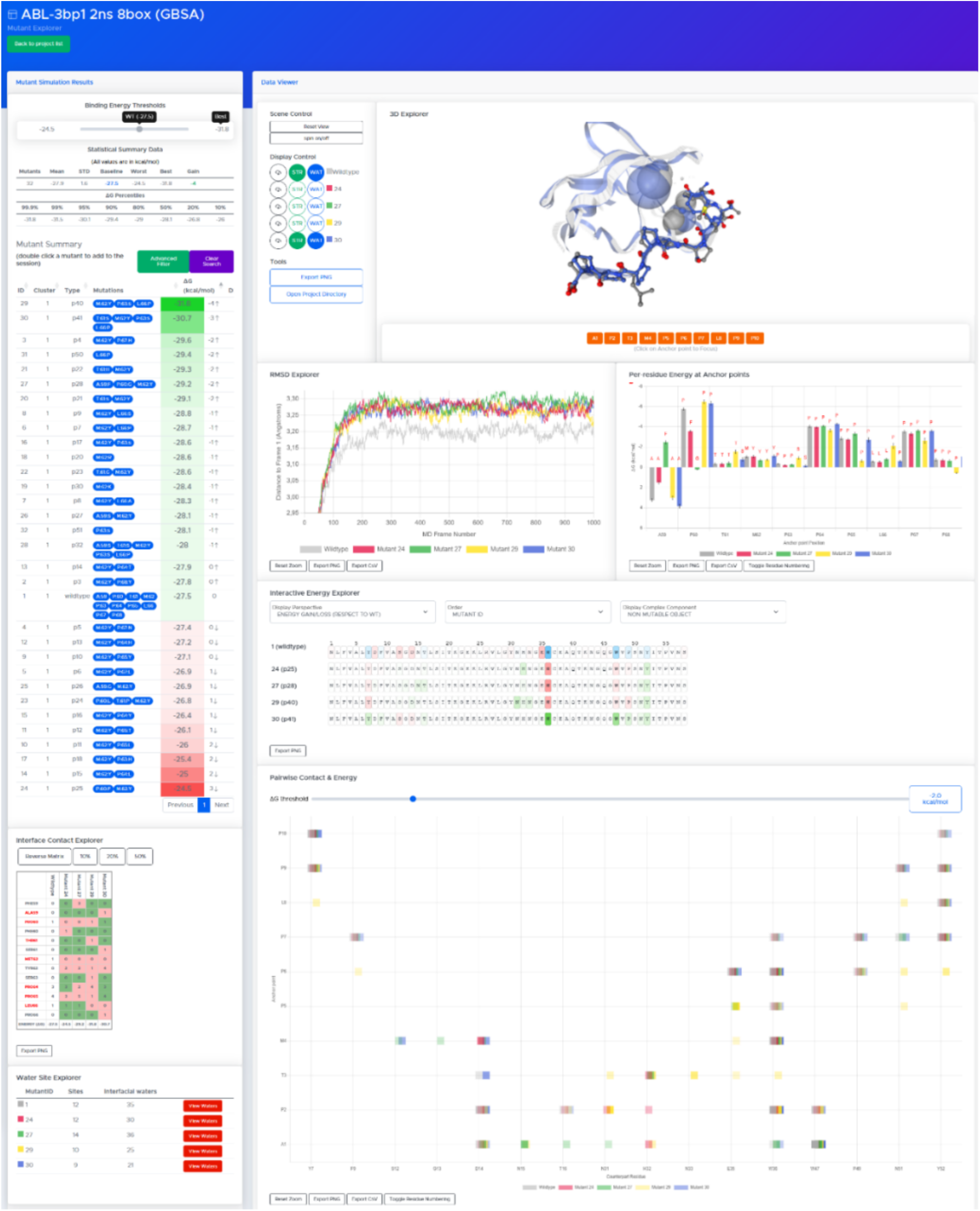
DyME’s TCA - Mutant Explorer tool. The “Mutant Simulation Results” panel is shown at the top left side. The Data Viewer panel and its widgets for exploration are shown at the right side and lower left area.

### Mutant Simulation Results Panel

In this panel, a set of widgets (*i.e.* Binding Energy Thresholds, Statistical Summary and Mutant Summary) provide a detailed report on the energetics of the complete mutant library, featuring an interactive tabular format that enables users to thoroughly analyze the energetics of the simulated mutants in comparison to the WT (Fig 9).

**Fig 9.**
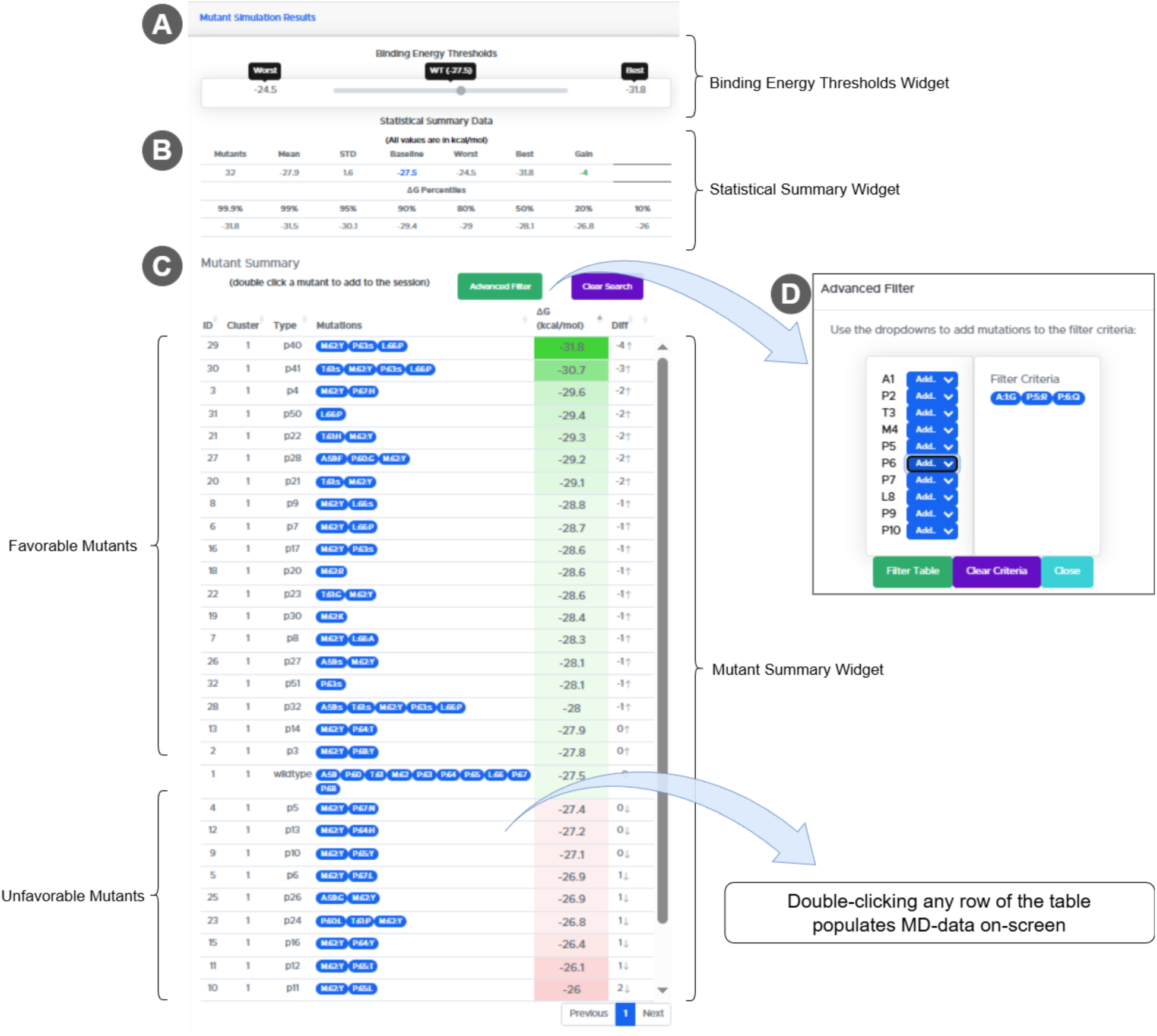
DyME’s TCA - Mutant Simulation Results panel. **(A) Binding Energy Thresholds widget**. Information is provided about the “worst” and “best” binding energy values in comparison to the WT system (labeled as “WT”). It offers a visual reference on how distant the baseline energy is with respect to that of the worst and best mutants. **(B) Statistical Summary Data widget.** This table provides the mean, the standard deviation, the baseline/worst /best energy values, as well as the maximum energy gain with respect to WT. It also displays energy percentiles (from 10% to 99.9%) summarizing the distribution of ΔG values among all mutants. **C) Mutant Summary widget**. This interactive table is built using the DataTables JQuery-plugin. Each row represents a particular mutant or WT system ranked by total energy values. Sortable columns provide information on mutant ID, cluster ID, type of mutant combination (*i.e.* singlet, doublet or triplet), mutations (*i.e.* position and residues involved), total ΔG in kcal/mol and the corresponding gain or loss with respect to the value obtained for the WT system. For analysis convenience, the widget table displays a green-to-red color gradient highlighting mutants from the most favorable to unfavorable energy compared to WT, respectively. **(D) Advanced Filter.** The mutant summary table can be dynamically filtered by combinations of point mutations using the button at its top. This filtering tool allows the user to define up to a triplet mutation as “filter criteria”, returning only the matching mutants that contain it. If no mutants were to meet the given criteria, the missing combination can be added to the mutant library and enqueued for simulation on-the-fly.

Each mutant, corresponding to a row in the “Mutant Summary” table (Fig 9C), is reactive to a “double-click” event that collects the corresponding data on that particular simulation and feeds the “*Data Viewer Panel*” (see below), which consists of several individual analysis widgets, including a 3D explorer for detailed visual analysis. In an explorative session, data can be loaded for as many mutants as desired and instantaneously populated on-screen.

### DataViewer Panel

All components under the *“Data Viewer”* panel (Fig 10 and 11) are context-aware and fully reactive. A crosstalk between the *"Mutant Summary”* table and the exploration session populates data asynchronously in the widgets using database-aggregated JSON objects. Each mutant added to the session is automatically assigned a particular color-code and a label. Clicking a particular mutant label at the base of each widget allows toggling that mutant On/Off. This action is propagated to all plot widgets showing or hiding information on that particular mutant in real-time. Furthermore, for analysis convenience, most widgets in the *“Data Viewer”* panel offer the option to export their content either as a high-resolution PNG image or a CSV file including the raw plot data.

**Fig 10.**
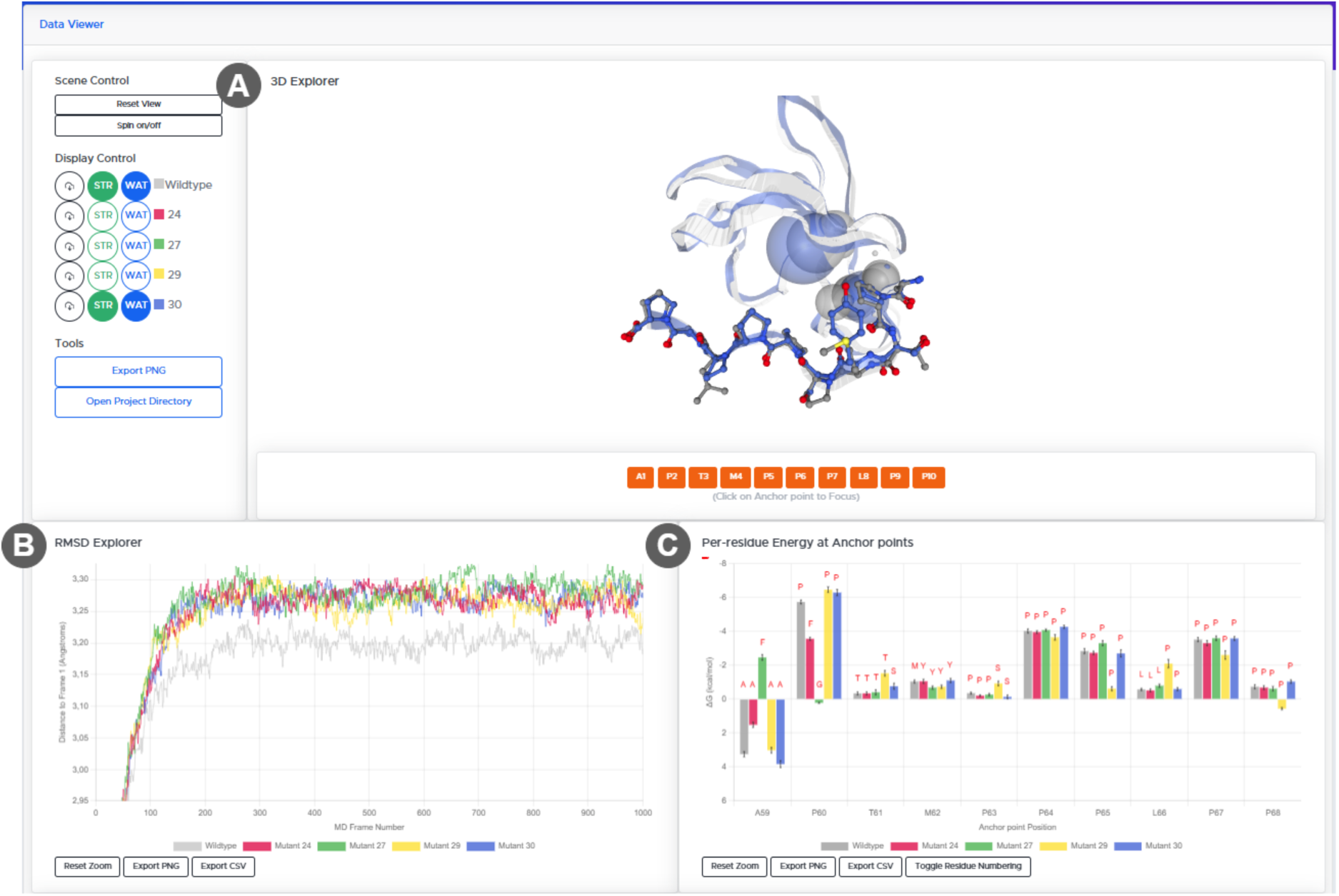
DyME’s TCA - DataViewer Panel 3D Explorer, RMSD Explorer and Per-residue Energy at Anchor-points widgets. **(A) 3D Explorer widget**. Built using NGLViewer [41,42], it displays the reference WT structure and automatically superposes the structure (best pose) of all mutants selected for comparison. Proteins are displayed as ribbons (with the corresponding mutant-color assigned), and anchor point side chains are shown in sticks. Interfacial water-sites are shown as spheres. The explorer allows molecular translation, rotation and zooming. At the left of the explorer window, the “Scene Control” offers reset-view and On/Off spinning options. The “Display Control” below allows the user to control individual viewing (On/Off) of each mutant (labeled by its ID and color code) and its corresponding water-sites by clicking the buttons labeled “STR” and “WAT”, respectively. Likewise, a button labeled with a “download icon” delivers the user a VMD [40] script (in TCL language) to display the MD trajectory with convenient predefined representations. This option also creates a VMD-compatible NetCDF [45] trajectory-file at the mutant’s output folder. **(B) RMSD Explorer widget.** Built using the ChartJS library, it displays a RMSD plot (*i.e.* MD-frame number in the *x-axis,* and distance to frame 1 in Angstroms in the *y-axis*) throughout the simulation time for any mutant selected for exploration. This plot is useful to inspect structural stability affected by the performed mutagenesis. The component embeds a responsive auto-refresh feature that adds or removes individual RMSD plots dynamically when selecting a new mutant for exploration. The mutant ID labels appearing at the bottom of the plot respond to the “mouse-click” event to toggle a mutant On/Off. A button at the bottom of the widget provides the means to reset the zoom (controlled through a wheel or touch mouse option). **(C) Per-residue Energy at Anchor Points widget.** Built using the ChartJS library, this graph provides a detailed comparative of the energetic effects of each mutation at each anchor point in a cumulative and fully interactive fashion. Anchor points (residue name and number) are represented in the *x*-axis, and their corresponding energy values (in kcal/mol) in the *y*-axis. Each bar is labeled at the top with the amino acid at that position and the corresponding error bar for the standard deviation of the energy value. Hovering the mouse on each bar shows its per-residue energy value (in kcal/mol) and standard deviation. Bars are colored by mutant ID, which allows to easily spot energy gains or losses. This is particularly useful when dealing with a large number of simulated mutants and trying to infer which recognition properties could be implemented in further engineering design strategies. For convenience (*i.e.* when a large number of mutants are in display), the plot can be zoomed as desired through a wheel or touch mouse option. A button at the bottom of the widget provides the means to reset the zoom. The *“Toggle Residue Number”* button offers the option to toggle residue numbering on-the-fly, which is convenient when the original PDB residue numbering has been altered. If such residue numbering mapping was provided during the preparation wizard (Fig 4C), DyME tracks the appropriate numbering conversions automatically.

**Fig 11.**
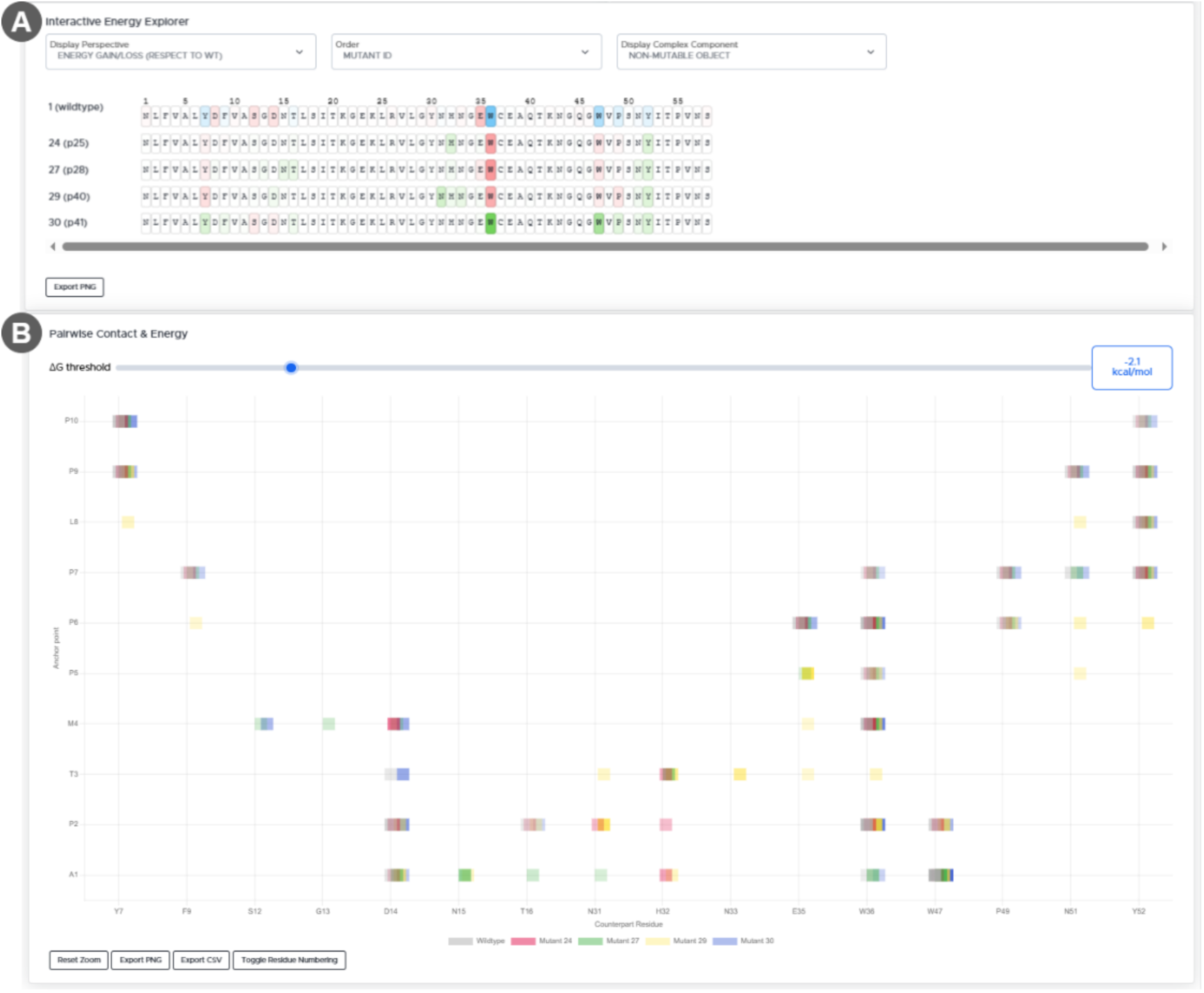
DyME’s TCA-DataViewer Panel Interactive Energy and Pairwise Contact Explorers. **(A) Interactive Energy Explorer widget**. It allows visual comparative exploration of high-content per-residue energetic data. The amino acid sequence of the systems under investigation (*i.e. “mutable”* and *“non-mutable”* objects) are displayed, and they are responsive to the mouse-hovering event over each position to display its per-residue energy value (in kcal/mol). At the top of the widget, in the *“Display Perspective”* drop-down, three distinct options are offered for energetic analysis: *i) Energy distribution*: For each mutant being analyzed. This view highlights residues that contribute most strongly to the total binding energy using colored boxes: blue and red for stabilizing and destabilizing contributions, respectively. The color intensity fades gradually to reflect the relative strength of each residue’s contribution. *<u>ii</u>*) *Energy gain or loss* (with respect to WT): This display option emphasizes positions with improved or worsened *per residue* energy with respect to WT. Energy variations are provided in a green-to-red gradient representing gain or loss with respect to the same position in WT, respectively. *iii) Standard deviation of energy (*at individual positions per MD run): This display option highlights the energetic variability at each position throughout the MD simulation, helping to reveal fluctuations over time. Color transparency indicates the relative magnitude of the energy values, with greater opacity and transparency corresponding to higher or lower values, respectively. **(B) Pairwise Contact & Energy widget.** Built using a modified version of the heat-map plugin of the ChartJS library, this graph offers an interactive map of all pairwise interactions (*i.e.* the contacting residues and the corresponding pairwise energy value) for all selected systems on-the-fly, and it allows interactive and selective analysis per mutant. The *y*-axis represents anchor point positions at the “mutable object”, and the *x*-axis represents interacting residues at the binding partner or “non-mutable” object. Squares following the mutant ID color code represent the inter-molecular contacts. Mutant additions to this viewer are cumulative. Mouse-hovering over a particular square shows the pairwise energy contribution at the corresponding residue pair for each mutant exhibiting that particular pair-interaction (only shown when the total energy contribution is <-0.4 kcal/mol). If the same contact is conserved in several mutants, the squares at such position will overlap with a slight offset to the right, revealing mutant-unique contacts with relative ease. A horizontal slider at the top of the widget canvas (labeled “ΔG Threshold”) allows the user to adjust the transparency of each square depending on whether its energy value falls below or above the specified threshold. The mutant ID labels appearing at the bottom of the plot respond to the “mouse-click” event to toggle a mutant On/Off. The zoom level can be controlled through a wheel or touch mouse option.

The “*Interactive Energy Explorer*” can display the energetic information from the point of view of the “mutable object” and reveal energetic correspondence at the binding partner or “non-mutable object” (option by default); or *vice versa*. It allows elucidates whether certain mutations are forming novel interaction sites at the counterpart molecule, providing valuable insights for subsequent engineering steps.

On the other hand, the *“Pairwise Contact & Energy”* widget makes non-overlapping squares readily distinguishable at a glance, enabling rapid and intuitive identification of mutant-specific interactions. This analysis tool is particularly useful, as it eliminates the need for manual inspection of large volumes of energy decomposition data across mutants. It reveals the energetic effects of mutations across multiple positions in the receptor counterpart, which would otherwise require substantial time and effort to obtain. Furthermore, the energy thresholding feature assists in the discrimination between the relative strength or weakness of atomic contacts among all variants and, therefore, it facilitates the straightforward identification of key residues involved in recognition across the evaluated systems.

The “*Interface Contact Explorer”* (Fig 12) provides a dynamic interactive table for exploring interfacial contacts across simulations, enabling straightforward and comprehensive comparative analysis of multiple mutants and the WT, and is particularly useful for projects involving large mutant libraries. This component eliminates the need for manual time-consuming comparison of interfacial contacts across multiple mutants.

**Fig 12.**
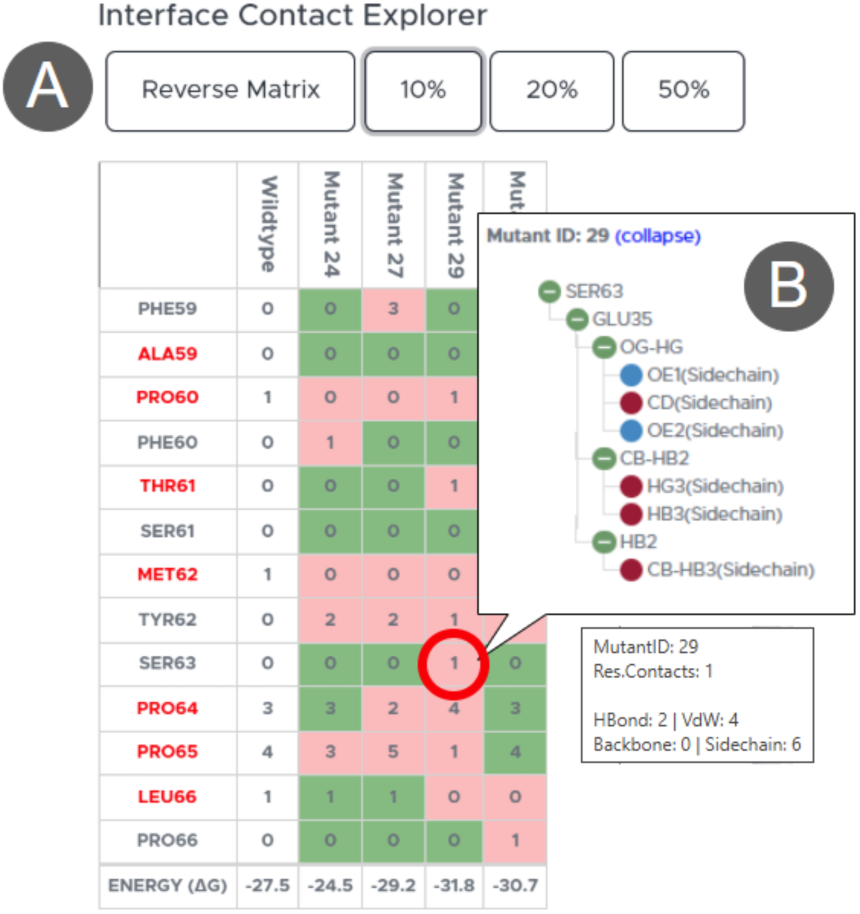
Interface Contact Explorer widget. **(A) Interface Contact Explorer**. A table containing the obtained number of contacts between the “mutable object” and the “non-mutable object” is built. Each mutant selected for comparison is rendered on a column and each row represents an amino acid position, which is labeled in red when it corresponds to an anchor point. By default, this table displays contacts from the perspective of the “mutable object” with a 10% threshold. The user can reverse the direction of the contact matrix (*i.e.* displaying contacts from “non-mutable” to “mutable object”) and display contact information at different frequency thresholds (*i.e.* 10%, 20% and 50%) by using the corresponding buttons at the top of the table. The bottom of this table shows the total binding energy (in *kcal/mol*) for each mutant. Each cell of the table shows the number of interfacial contacts and is colored in green or red depending on whether the number of contacts with respect to WT remains unchanged or not, respectively. Hovering over a cell displays a tooltip with the number and type of contacts. **(B)** A single-click on a particular cell of the table (as shown in **A**) deploys a collapsible list of the residues and atoms involved in the analyzed interaction, which are classified as “side chain” or “backbone” contact, color-coded in blue for polar and brown for hydrophobic.

### Water Site Explorer

The “*Water Site Explorer*” widget (Fig 13A) has been designed to provide in-depth information on interfacial water-sites with a simple mouse-click. Here, the user can analyze water-mediated interactions with the interactive exploration capabilities the *“Water-Contacts Summary”* panel (Fig 13B) and the 3D explorer (Fig 13C).

**Fig 13.**
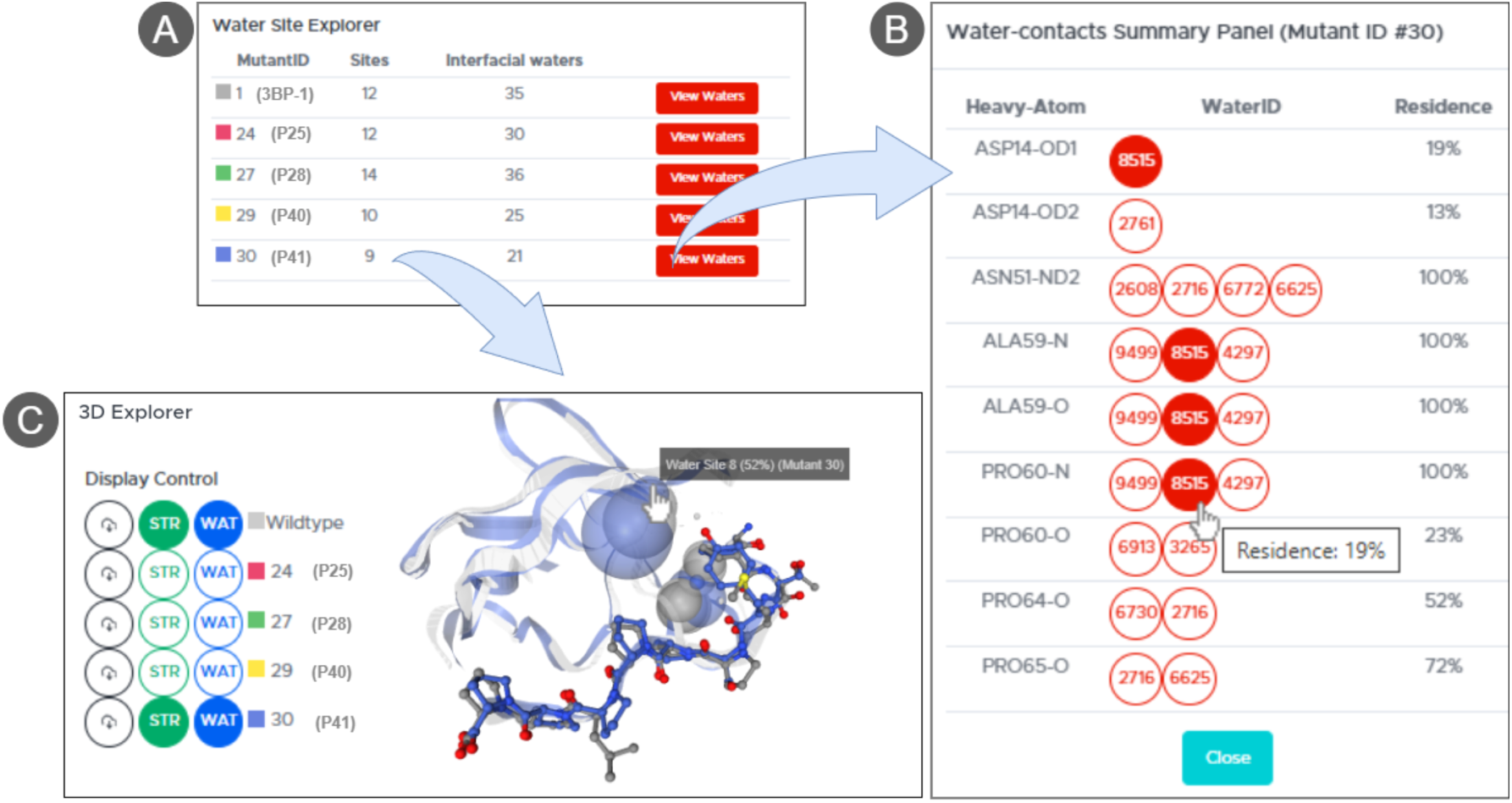
Components of the Water Site Explorer Widget and crosstalk with the 3D Explorer. **(A) Water Site Explorer**. It displays a table summarizing the number of water-sites and water molecules residing in them. **(B) Water-contacts Summary.** Clicking the red buttons labeled as “View Waters” in (A), a fully interactive table showing the atoms at the mutable and “non-mutable” objects involved in interactions with those waters is shown. A full listing of interfacial water-contacts is provided organized by residue (*i.e.* residue name, number and heavy-atom) and the ID of each involved water molecule, as well as the combined residence of all water molecules participating in that interaction across the simulation. The residence of a particular water molecule will appear by positioning the cursor on the ID icon. **(C) 3D Explorer widget**. Interfacial water-sites for WT and any selected mutant system are graphically shown as spheres, each representing the centroid coordinates of a water-site and sized proportionally to the combined residence time of its water molecules during the MD simulation.

This widget enables rapid identification of the involvement of interfacial waters and their interactions. Hovering the cursor over any water ID icon in the summary panel interactively highlights all instances of its interactions with other residues, providing a comprehensive visualization of water contacts at a single click. Likewise, the centroid representation provides intuitive clues on the emergence, disappearance or conservation of water-sites across mutants.

### Specificity Finder

The “*Specificity Finder”* tool (Fig 14) streamlines the analysis of differential binding energies between two projects that share common ligand mutants in their libraries but comprise different protein receptors. Thus, this module allows a straightforward comparison of two DyME projects simultaneously, which is particularly helpful for investigating specificity (*i.e.* identifying mutations that increase affinity for one receptor while simultaneously decreasing it for another). By leveraging database aggregations and scavenged energy decomposition records, this tool generates a list of up to 50 mutants common to both projects, thus providing the user with a powerful view to infer insights for improved specificity. As shown in the provided example in Fig 14, the first row of the table, mutant ID 29 represents a triplet mutation (M4Y, P5S, L8P) that increases specificity for the protein receptor in project ABL1-3bp1 (from -27.5 to -31.8 kcal/mol) and simultaneously decreases it for the protein receptor in project FYN-3bp1 (from -26 to -16.8 kcal/mol).

**Fig 14.**
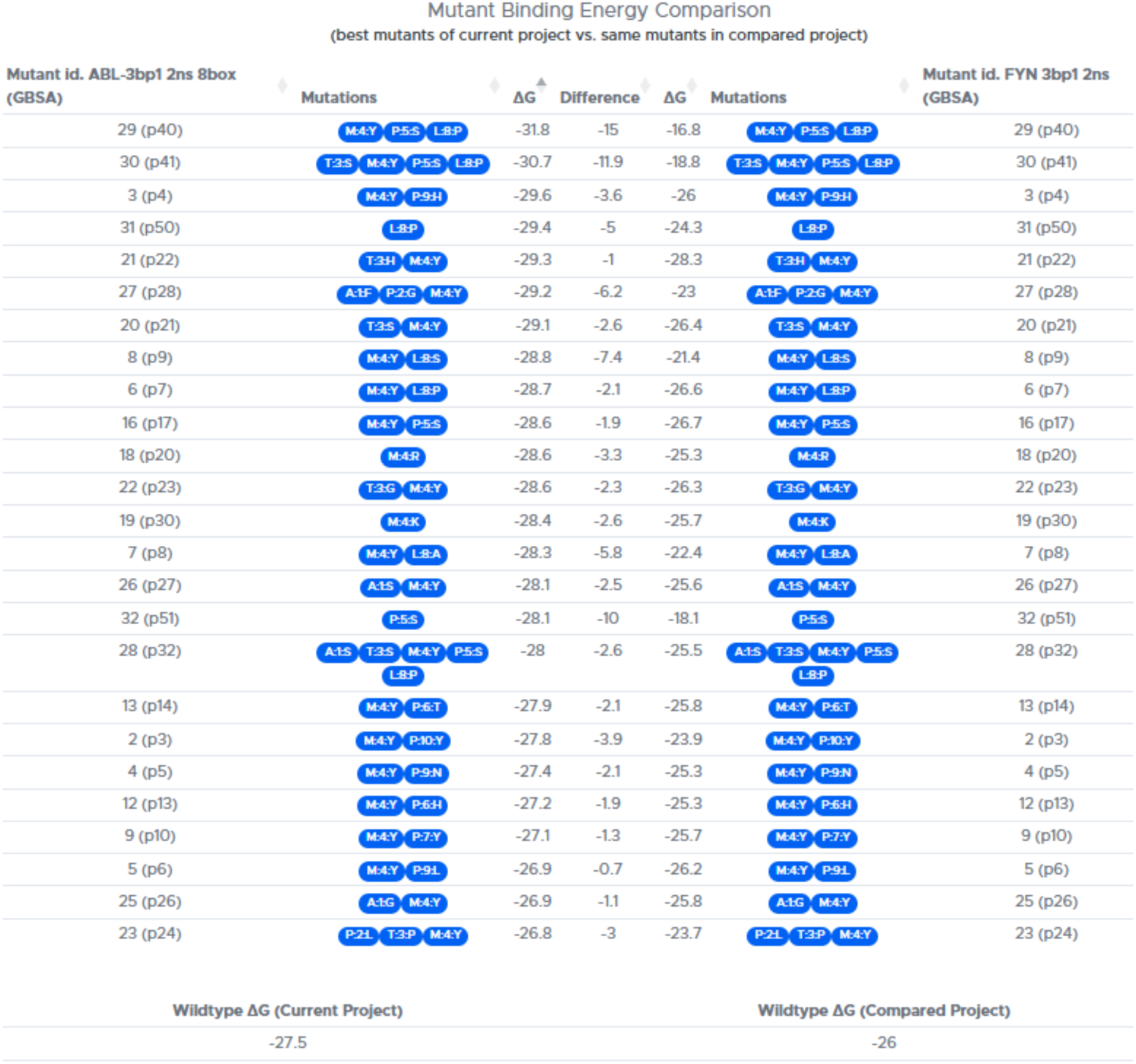
Specificity Finder. Comparative example of two projects sharing the same mutable object (*i.e.* a peptidic ligand) but different non-mutable objects (*i.e.* a protein receptor).

### Case Study

We have used experimental data from a previously reported study on protein mimicry engineering to test DyME’s utility in guiding a design rationale for new protein ligands. SH3 (SRC Homology 3) are small protein domains [46] known to regulate cell signaling processes through interactions with poly-proline II (PPII) motifs from partner proteins. Such interactions are mostly characterized by hydrophobic contacts of the prolines at an exposed hydrophobic patch on the SH3 domain surface. Additionally, water-mediated interactions have been reported to contribute to binding [38]. A major challenge in engineering effective mimetics of SH3 binders is the high promiscuity of their interactions and the low affinity exhibited by natural target sequences. For instance, the SH3 domains of the Abl and Fyn tyrosine kinases bind with the same low affinity the peptide 3bp-1 (APTMPPPLPP*, K*_d_ = 34 μM) corresponding to the sequence of their first identified native target [47]. The sequence of 3bp-1 was systematically mutated to generate variants of this SH3 recognition motif with improved binding affinity and specificity, allowing discrimination between Abl and Fyn SH3 domains. Using molecular modeling and detailed structure-based analysis, the interactions of the mutants with both protein receptors were evaluated *in silico*, leading to the design of the peptide p41 (APSYSPPPPP), which experimentally showed a 20-fold increase in binding affinity for Abl-SH3 and a 10-fold decrease for Fyn-SH3 [36].

We have tested DyME’s utility by replicating the peptide ligand mutagenesis rationale used by the authors in this study. For this, we defined all positions of the 3bp-1 peptide as *anchor points* for HTP mutagenesis and analyzed the impact in recognition by the two protein receptors using DyME. We showcase how the analysis of the obtained MD results is made easy and straightforward by DyME. Details on input file preparation, simulation parameters, anchor point selection rationale, generation of mutant structures, additional calculations and in-depth use of DyME for analysis are provided in the Case Study chapter on S3 Appendix. The generated inputs, MD simulation outputs and raw outputs for this case study have been deposited at Zenodo [48] and are available for download at https://zenodo.org/records/18014320.

### Availability and future directions

DyME represents an open-source, scalable platform that integrates high-throughput mutagenesis and automated MD simulations, enabling systematic investigations of protein recognition mimicry at large-scale. Its feature-rich analysis toolbox provides a highly interactive environment for comparative exploration of energetic and molecular recognition features across a large collection of ligand/receptor systems.

DyME can effectively guide rational design for affinity and selectivity optimization by enabling straightforward exploration of broad mutational spaces through high-content MD simulation data. Its interactive analysis environment allows systematic analysis of mutational effects and complex interfacial interaction patterns, thereby facilitating more informed and efficient biomolecular engineering.

DyME’s distributed architecture makes it easily scalable and adaptable to existing computational infrastructures. Its design combines modern software design paradigms with a document-based database, providing efficient access to information and seamless orchestration of its components, while addressing critical bottlenecks traditionally associated with large-scale MD-based studies, otherwise labor-intensive, error-prone, and challenging to attain. By overcoming the limitations of conventional fragmented workflows, DyME facilitates more efficient large-scale systematic MD-based investigations.

DyME has the potential to become a cornerstone platform for accelerated rational design, integrating large-scale MD simulations and machine learning to rapidly identify and predict exploitable protein recognition features.

## Acknowledgements

We are grateful to Prof. Frank Buchholz, Martina Augsburg and Dr. Liliya Mukhametzyanova for support and helpful discussions during the development of the protein-DNA feature of DyME. We also acknowledge Dr. Peter Eastman, Dr. Jason Swails, Dr. John Chodera, Dr. Ben Webb, Dr. Alexander Rose and their teams for their valuable assistance with inquiries regarding OpenMM, MDTraj, Modeller and NGLViewer.

The authors acknowledge high-performance computing time at the NHR Center of the TU Dresden, which is jointly supported by the Federal Ministry of Education and Research and the state governments participating in the NHR (www.nhr-verein.de/unsere-partner).

## Supporting Information

**S1 Appendix. Software Dependencies.** Contains a table with third-party libraries and dependencies, technical details on GPU requirements, and an overview of the database structure.

**S2 Appendix. Details on Methods Implementation**. Contains additional details on implementing methods for automatic mutant library generation and database storage, as well as other methods of the workflow.

**S3 Appendix. Case Study**. Contains detailed information on input generation and the creation of projects for the case study, as well as in-depth comparative analysis examples using the DyME TCA.

**S4 Appendix. Installation and MD Simulation test-data**. Contains instructions on how to locate, understand and install the DyME test data.

